# Maximising imaging volumes of expanded tissues for inverted fluorescence microscopy

**DOI:** 10.1101/2025.09.12.675899

**Authors:** Miguel Cardoso Mestre, Jacob R. Lamb, Madeline A. Lancaster, James D. Manton

## Abstract

Expansion microscopy (ExM) has enabled nanoscale imaging of tissues by physically enlarging biological samples in a swellable hydrogel. However, the increased sample size and water-based environment pose challenges for deep imaging using conventional inverted confocal microscopes, particularly due to the limited working distance of high-numerical aperture (NA) water immersion objectives. Here, we introduce a practical imaging alternative that utilizes an inverted water-dipping objective and a refractive-index-matched optical path using fluorinated ethylene propylene (FEP) film. Through point spread function (PSF) measurements and simulations, we show that the FEP film introduces predominantly defocus-like wavefront profiles characteristic of high NA systems, which result in an easily correctable axial shift of the focal plane. To ensure stable immersion and refractive index continuity, we use an arrangement relying on an FEP film, Immersol W, water and a FEP-based imaging dish. This configuration achieves sub-micron lateral and axial resolution, supports large tile-scan acquisitions, and maintains image quality across depths exceeding 800 *µ*m. We validate the system by imaging 4×-expanded U2OS cells and human cerebral organoids. Our approach provides a low-cost, plug-and-play solution for high-resolution volumetric imaging of expanded samples using standard inverted microscopes.

## 1 Introduction

Since its introduction in 2015, expansion microscopy (ExM) has emerged as a powerful method for improving spatial resolution in biological imaging by physically enlarging hydrogel-embedded samples [1]. By isotropically expanding the sample, ExM enables conventional light microscopes to resolve nanoscale structures without the need for complex super-resolution instrumentation. The method has been broadly adopted in diverse biological contexts, including studies of cells and tissues in their native environments [2–4]. More recently, Tavakoli *et al*. demonstrated the potential of ExM for light microscopy-based connectomics [5]. A key advantage of ExM is that the expansion process reduces light scattering by homogenizing the refractive index of the sample to that of water (n = 1.333), thereby effectively clearing the tissue. This dual benefit of enhanced resolution and improved optical transparency makes ExM especially well-suited for imaging thick biological specimens at high spatial resolution across large volumes.

Due to the aqueous nature of the sample, water immersion objectives are the standard choice for imaging. However, the inherent trade-off between numerical aperture and working distance for coverslip-corrected lenses means that standard high-numerical aperture (NA) water immersion objectives typically have short working distances, often less than 300 µm. This presents a significant challenge when imaging along the optical axis, particularly for thick samples, as the physical expansion of the sample increases its overall dimensions. This makes it impossible to access deeper regions in standard inverted confocal microscopes. Indeed, for 4*×* expansion, less than 75 microns of depth can be imaged. As a result, imaging of entire expanded tissue samples requires a strategy that accommodate both the expanded sample size and its water-based environment.

Several research groups have proposed novel gel chemistries that increase the stiffness of the hydrogel, making it more amenable to mechanical sectioning. The Zeng group introduced a protocol for Expansion Tomography, reporting a gel formulation with more than twice the elastic modulus than that of other popular methodologies such as ExM, MAP, or U-ExM [6]. This increased rigidity was achieved by incorporating AMPS-Na (2-acrylamido-2-methyl-1-propanesulfonic acid) into the gel recipe and fine-tuning the concentration of N,N’-methylenebisacrylamide to enhance mechanical strength without compromising the expansion factor. More recently, in 2024, a new formulation was introduced in which sodium acrylate was replaced by potassium acrylate [7]. This substitution resulted in a stiffer gel, which may further facilitate mechanical sectioning due to its increased mechanical robustness. These approaches aim to enable volumetric reconstruction of the entire sample through serial imaging of mechanically sectioned expanded tissue. However, mechanical sectioning remains a labour-intensive, error-prone process that requires specialised and costly equipment, making it less than ideal for many applications, especially as images of separate sections must be computationally aligned to form a whole tissue volume.

As an alternative, several custom microscopy platforms have been developed to image entire expanded tissue samples, including ExA-SPIM and Curved Light Sheet systems [8, 9]. These systems offer large fields of view (FOVs), exceeding 10 mm, and can bypass the working distance limitations of conventional high-NA objectives. However, this increase in FOV typically comes at the expense of spatial resolution. While resolution can be partially recovered by selecting ExM protocols with higher expansion factors, achieving the resolution equivalent to 4*×* expansion in a confocal microscope may require iterative protocols exceeding 20*×*. In such cases, imaging with a conventional confocal microscope may be more practical. Additionally, the construction and alignment of such systems requires specialized knowledge in optical physics and access to expensive components, which may limit their accessibility in many laboratories.

As an alternative to current imaging approaches for large, expanded samples, we present a practical framework that extends imageable depth without compromising resolution, and is fully compatible with standard inverted microscopes. Our approach makes use of a water-dipping objective (in this demonstration a Nikon CFI Apo NIR 60X W) mounted in an inverted orientation to take advantage of its extended working distance (2.80 mm vs 0.28 mm for the corresponding 60*×* water-immersion objective). Water-dipping objectives are designed to operate with direct immersion in aqueous environments and are highly sensitive to mismatches in the optical path, which can introduce spherical aberrations and degrade resolution. To preserve their imaging performance, we ensured that the entire optical path between the objective and the sample is closely matched to the refractive index of water. This was achieved by employing a custom imaging dish fitted with a 127 µm-thick fluorinated ethylene propylene (FEP) film, as used with filament-extrusion 3D printers. The refractive index of FEP (*n =* 1.344) closely matches that of water, allowing effective refractive index matching with negligible spherical aberration.

In this manuscript, we detail the complete imaging configuration, including our approach for refractive index matching between the sample and the objective. For our demonstration lens, this approach achieves a measured resolution of 0.392 *±* 0.040 µm *×* 0.396 *±* 0.040 µm *×* 1.744 *±* 0.235 µm, ignoring the expansion factor. We perform point spread function (PSF) simulations to assess the impact of the use of the FEP film in imaging quality, evaluating the introduction of optical aberrations. We demonstrate the performance of the framework by imaging 4*×*-expanded U2OS cells and human cerebral organoids, acquiring an image composed of 600 tiles. The increase in imageable depth obtained by the use of a water dipping lens was compared to a conventional water-immersion objective (CFI Plan Apo VC 60XC WI). Finally, the resolution as a function of depth was also quantified, confirming that imaging performance is preserved with depth, validating the framework’s robustness for increasing the imageable axial depth for expanded samples.

## 2 Results

Water-dipping objectives have much larger working distances than water-immersion lenses, but are not corrected to image through a coverglass. As such, these lenses are conventionally not suitable for use on inverted microscopes, i.e. the predominant form of high-resolution fluorescence microscope found in microscopy facilities and research laboratories. To mitigate aberrations introduced by imaging through a conventional glass coverslip, we explored the use of a commercially available 127 µm-thick fluorinated ethylene propylene (FEP) film as the base of an imaging dish (A7816: Attofluor™ Cell Chamber, ThermoFisher Scientific). Due to the close match between the refractive index of FEP (*n =* 1.344) and water (*n =* 1.333), we hypothesised that this substitution would reduce refractive index discontinuities and thereby minimise optical aberrations.

To test this hypothesis, we decided to compare the theoretical imaging performance of a 1.0 NA water dipping lens when used with either an FEP or glass coverslip. To this end, we calculated the pupil aberration function, and simulated the corresponding 3D point spread functions (PSFs) using custom MATLAB code. These calculations were based the equations previously described by Booth et al. [10].

In particular, for a normalised pupil coordinate *ρ*, a maximum collection angle for the lens of *α*, a maximum collection angle when imaging in the coverslip medium *β*, and a coverslip thickness *d*, the phase aberration function is given by:

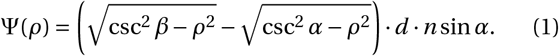

We evaluated this function for a 170 µm-thick glass coverslip (*n =* 1.52) and a 127 µm-thick FEP film (*n =* 1.344). The resulting phase aberration functions were plotted as a function of the normalised pupil coordinate (Figure 1a). While the glass coverslip introduces larger aberrations compared to FEP, the shape of the functions suggested that much of the profile can be described as high-NA defocus. To verify this, we computed a high-NA-defocus phase function for each of the coverslips and performed a search for the amount of defocus that minimised the overall root mean square error. The optimal correction required a phase shift of 1.3342 waves for FEP and 24.2713 waves for glass.

**Figure 1:**
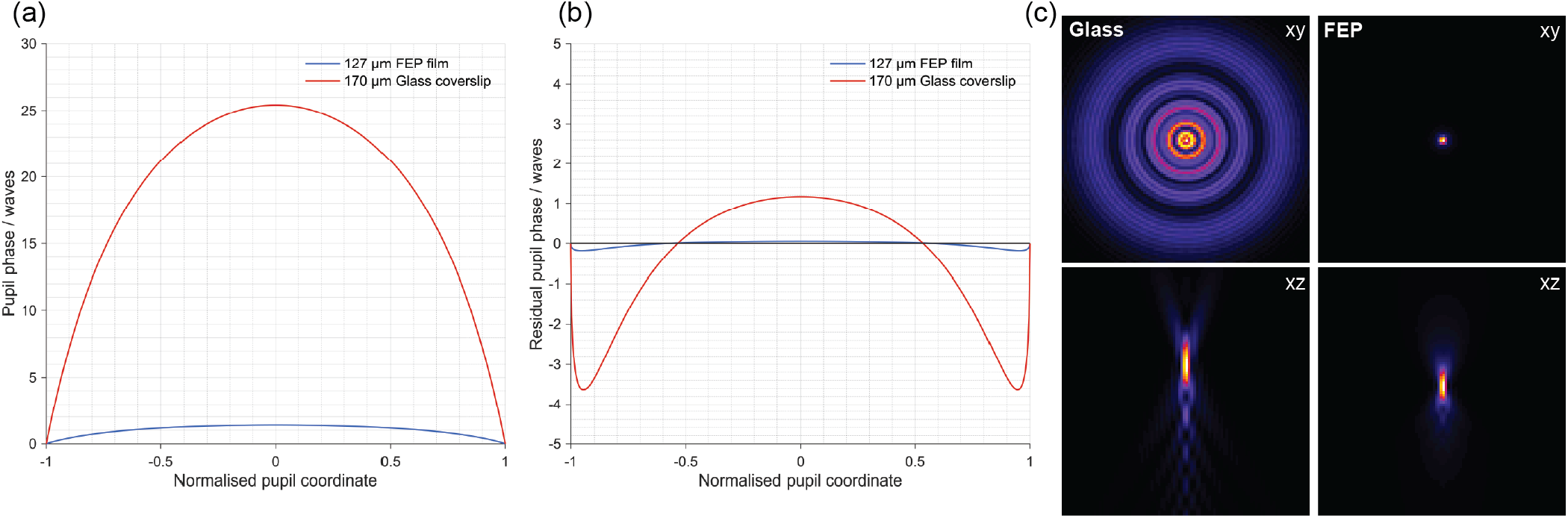
(a) Simulated phase aberrations when imaging through a 127 *µ*m FEP or 170 *µ*m glass coverslip, presenting a profile dominated mainly by full-defocus. (b) Residual aberrations after full-defocus correction, showing near-complete correction for FEP but not for glass. (c) Simulated 3D PSFs after full-defocus correction, with glass producing an aberrated PSF while the FEP coverslip preserves a near-ideal PSF.

Figure 1b shows the residual aberration functions,*ϕ*_res_ *= ϕ*_aberration_ *− ϕ*_full defocus_, for both coverslip types after full-defocus correction. For FEP, the residual aberration was minimal (less than 0.2 waves peak-to-valley), confirming that full defocus correction effectively restores the wavefront. In contrast, significant aberrations remained across the entire pupil when using the glass coverslip, indicating that additional corrections would be required. The residual aberration function was then used to compute the pupil function of the system, using the apodisation correction 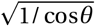 to account for the high numerical aperture (NA) of the objective lens being used.

3D point spread functions (PSFs) were then computed from the pupil function. Figure 1c shows the full-defocus corrected 3D PSFs for both coverslips. The results suggest that full-defocus correction sufficiently compensates for the FEP film’s refractive index mismatch with water, removing the need for further optical components. The PSF simulated when imaging through glass, however, presents noticeable spherical aberrations. Given that imaging through a glass coverslip would require more complex correction and we aimed to establish a plug-and-play framework for inverted microscopes, all subsequent experiments were conducted using the FEP film.

However, the use of an appropriate coverslip is not the only factor that requires consideration, and the selection of immersion medium also affects imaging quality. While water would be a natural choice, practical issues arose due to the long working distance (2.8 mm) of the objective. We found that a water droplet placed directly on top of the lens would not consistently maintain contact with the FEP film at shallower imaging depths (i.e. where the column of water required is at its most extended), resulting in degraded image quality.

As an alternative, we attempted to use an eye gel lubricant as an immersion medium, as its use has previously been reported for imaging applications[11] (Figure 2a). This type of gel has a refractive index close to water (1.338) and higher viscosity, offering greater stability. To assess its performance, we imaged 200 nm TetraSpeck microspheres dried on FEP film under widefield conditions. The acquired PSFs presented strong spherical aberrations (Figure 2b), indicating that this gel is not an effective immersion medium for high resolution imaging with a water-dipping lens. The aberration introduced into the PSF was also noticeable when imaging 4*×*-expanded U2OS cells labelled for microtubules and the nucleus (Figure 2c).

**Figure 2:**
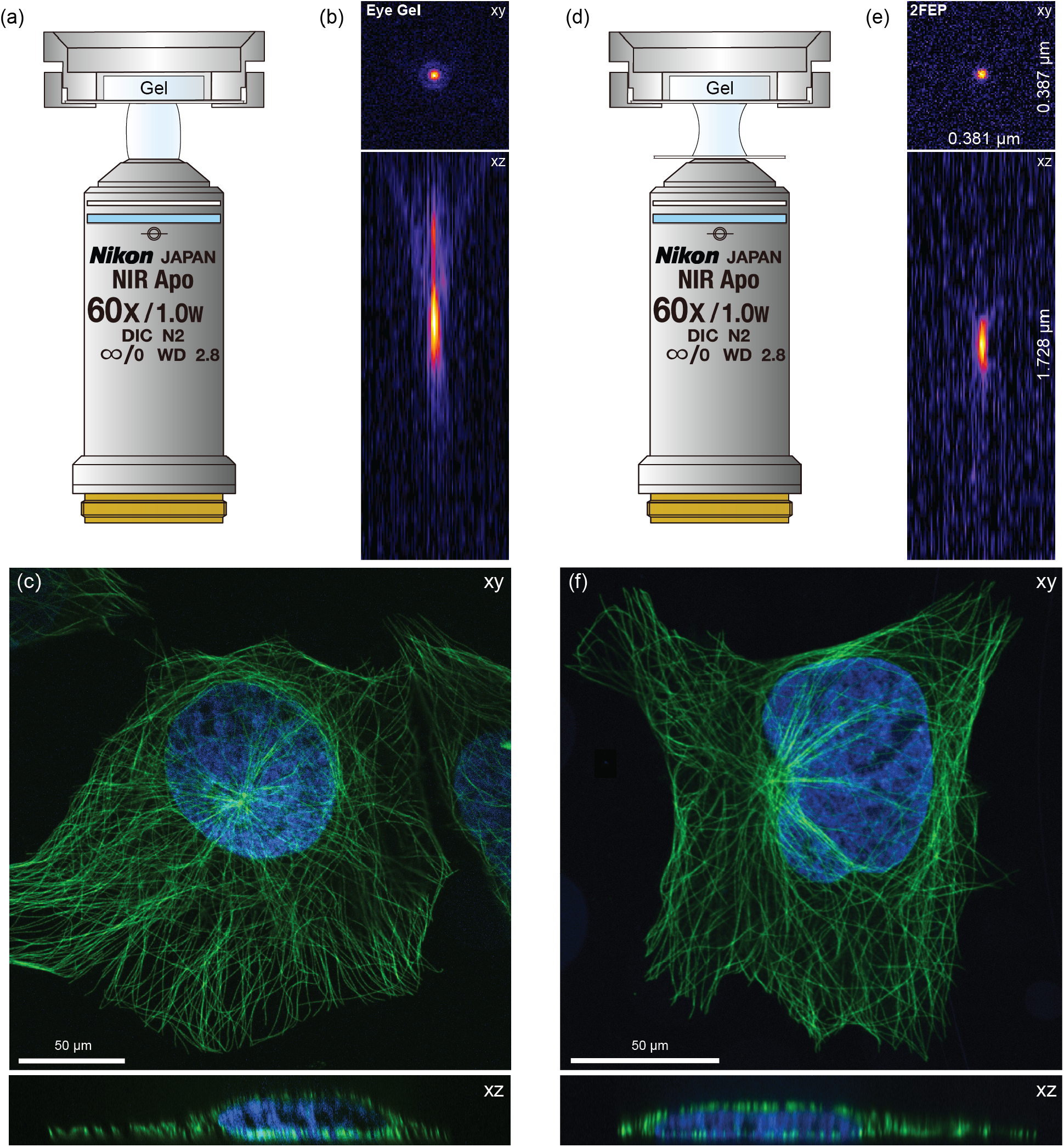
(a) Schematic of the imaging configuration using eye gel as the immersion medium, with (b) the corresponding experimental PSF showing a distorted, aberrated profile. (c) U2OS cells labeled for microtubules (*α*-tubulin) imaged with eye gel immersion, showing degraded image quality. (d) Schematic of the configuration using two stacked FEP films, with (e) the corresponding PSF exhibiting near-ideal lateral and axial profiles. (f) U2OS cells labeled for microtubules (*α*-tubulin) imaged with water immersion and two FEP films, showing high image quality.

To enable the use of water as the immersion medium, we considered placing a second FEP film resting on top the objective (Figure 2d). As FEP is more hydrophobic than glass, we theorised that this would alter the contact angle of the water droplet to one more suitable for maintaining an elongated column of water. To verify that the introduction of a new FEP interface would not hinder imaging performance, the appropriate PSF was both simulated and measured (Figure 2e, Supplementary Figure 1). For the purpose of the simulations, the FEP coverslip thickness was considered to be double that of previously, to account for the two films in the optical path. It was possible to visualise that once again the phase aberration function could be corrected effectively through full defocus alone. The pupil phase shift correction was estimated to be 2.67 waves. Imaging the same U2Os expanded cells as before now showed no noticeable aberrations (Figure 2f), particularly in the XZ view. Furthermore, imaging 4*×*-expanded HeLa cells labelled for the nuclear pore complex components NUP96 and NUP98 with either a glass coverslip and the two layers of FEP shows a much cleaner image for the latter condition (Supplementary Figure 2).

Similarly to the case of the eye gel, we also imaged 200 nm fluorescent beads to evaluate the point spread function (PSF). The achievable resolution was determined by fitting a 3D Gaussian function to the acquired bead data. The full width at half-maximum (FWHM) was computed for each axis from the fitted standard deviation, *σ*, where 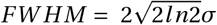. Analysis of 9 beads yielded a mean resolution of 0.392 *±* 0.040 µm *×* 0.396 *±* 0.040 µm *×* 1.744 *±* 0.235 µm, consistent with simulations and confirming the effectiveness of the immersion approach.

In practice, we found that while this arrangement worked well for imaging a small lateral extent, the FEP film placed onto the objective could sometimes become dislodged, particularly when acquiring large tilescans. To obviate this issue, we replaced the water between the bottom FEP film and the objective with Immersol W as, while this has the same refractive index as water, is much more viscous. We also reduced the size of the bottom FEP film to half the size of the top film (12.5 mm vs 25 mm, Supplementary Figure 3). With this, PSF measurements showed no degredation of the resolution from the fully-water case.

To evaluate the increase in usable imaging depth, we imaged expanded cerebral organoids stained with a pan-protein stain using NHS-ester [12]. Using the water dipping lens, the entirety of the depth of the organoid (1300 µm) could be imaged (Figure 3a). In contrast, using a water immersion objective (CFI Plan Apo VC 60XC WI), only the first 280 µm of depth could be imaged (Figure 3b). To further evaluate that lateral resolution was preserved at depth, we computed the mean tile Fourier-ring correlation (mtFRC) as a function of depth for a z stack acquired at the centre of an expanded organoid [13]. We chose to label this organoid for the transcription factor SOX2 as this would give a punctate staining, ensuring the presence of high spatial frequencies in the images. As shown in Figure 3c, resolution remained approximately constant over the first 800 µm in depth. Beyond this, we attribute the steady deterioration of resolution as a function of depth to pinhole crosstalk, a common issue when imaging using spinning disk microscopes at higher depths. Unfortunately, the spinning disk system used did not have the option to change to a disk with more widely spaced pinholes to counter this, although such disks are commercially available. Finally, we tested the robustness of the FEP-water-FEP-Immersol W construction by performing a large-scale multi-tile acquisition comprised of 600 tiles (Figure 3d), with a comparison between a random subset of tiles showing consistent imaging quality (Figure 3e–g).

**Figure 3:**
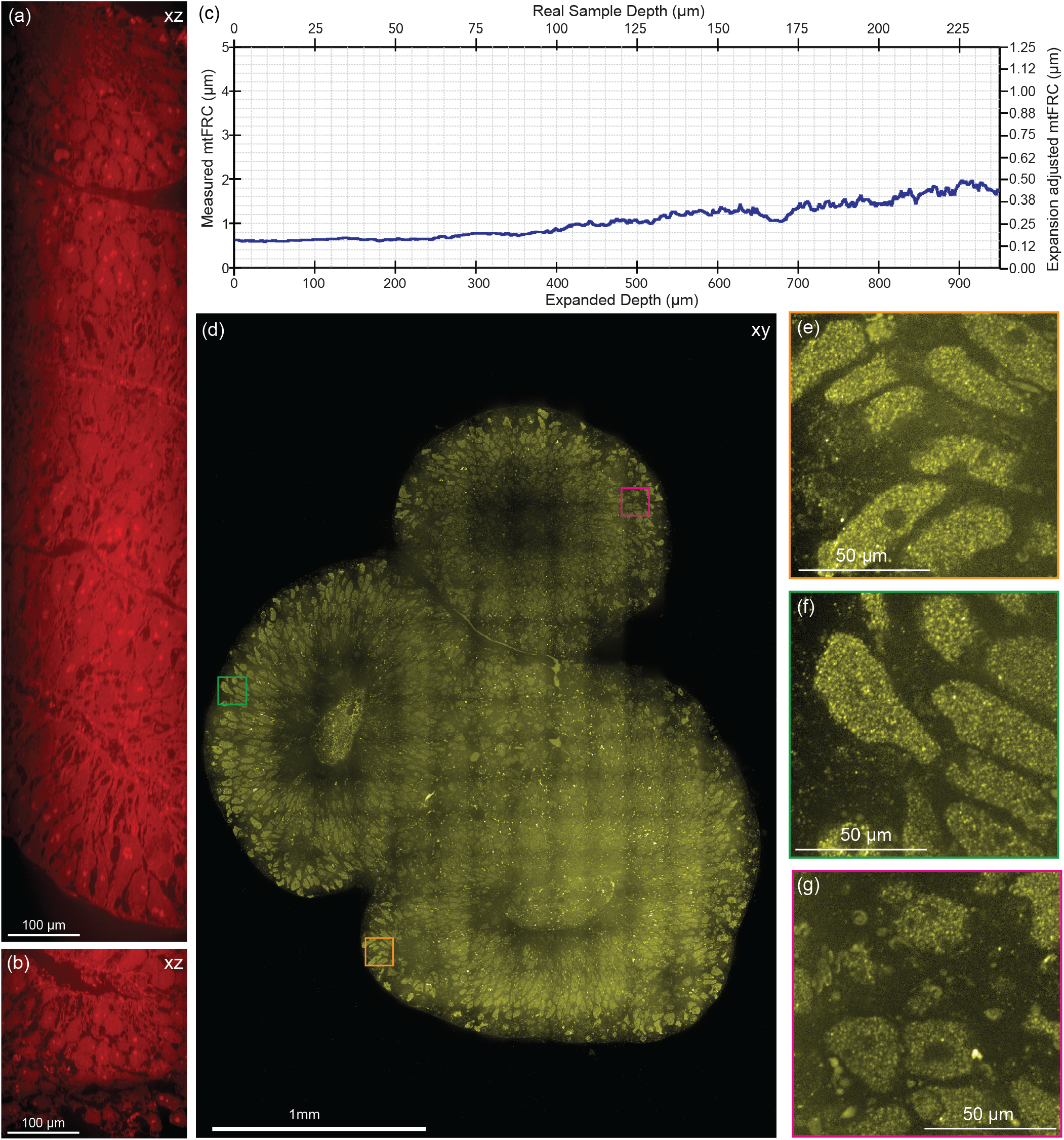
Comparison of imaging pan-stained expanded cerebral organoids using (a) a CFI Apo NIR 60× water-dipping objective and (b) a CFI Plan Apo VC 60× water-immersion objective, showing that the water-dipping approach markedly increases the accessible imaging depth. (c) mtFRC analysis demonstrates consistent resolution up to 800 *µ*m. (d) Large-area multi-tile acquisition of 4*×*-expanded cerebral organoids immunolabeled for the transcription factor SOX2. (e–g) Magnified views of the regions highlighted in (d).

## 3 Discussion

Here we present a simple and cost-effective approach for significantly extending the imageable depth of expansion microscopy (ExM) samples using a commercial inverted microscope. Our method uses a water-dipping objective in an inverted configuration, taking advantage of its extended working distance through a minimalistic setup that includes two 127 µm-thick FEP coverslips, water, Immersol W, and an AttoFluor chamber.

Although the inclusion of FEP film in the optical path introduces aberrations due to the refractive index mismatch, our simulations demonstrate that the resulting aberration profile is similar to that of a high-NA defocus and so can be easily compensated for by changing the focus of the microscope objective. Crucially, this aberration is not depth-dependent and so merely bringing the bottom of the sample into focus is enough to ensure unaberrated imaging throughout the volume.

Unlike other solutions that depend on complex and high-cost instrumentation, all our required equipment can be purchased for less than 5000 Euros, with the main expense being the water dipping lens. This ensures wide applicability and ease of integration into existing imaging workflows, facilitating volumetric imaging of large expanded tissues across a broad range of laboratories. However, while enabling significantly deeper imaging compared to standard water immersion objectives, the total imageable depth remains ultimately constrained by the working distance of the water-dipping lens of choice. Hence, the choice of objective should be carefully considered to ensure that the whole tissue thickness is accessible when imaging. As a result, one may need to select a lower NA lens, compromising the achievable resolution, in order to increase the imageable axial depth. In situations where this is not possible, or where the expanded sample exceeds the axial range of such objectives, alternative strategies such as mechanical sectioning with stiffer gel chemistries may still be required. Nevertheless, our approach offers a more accessible and less error-prone alternative to mechanical sectioning or custom-built light-sheet systems, applicable for medium sized samples, and can be combined with sectioning to image even larger samples.

## 4 Methods

### 4.1 Mammalian cell culture

Human U2OS osteosarcoma cells (ATCC) and HeLa FlipIn cells (gift from Simon Bullock) were cultured in DMEM-Glutamax (Gibco) supplemented with 10% fetal bovine serum (FBS, Gibco) at 37°C with 5% CO2. In preparation for expansion, cells were seeded onto 6 mm coverslips (Agar Scientific) and incubated overnight. In the morning cells were fixed for 30 minutes using 4% PFA and washed with PBS 3 times for 15 minutes.

### 4.2 Stem cell culture

A hESCs cell line (H9, female) was used to generate cerebral organoids and was cultured in feeder free conditions. Cells were grown in StemFlex (Thermo Fisher, A3349401) on Matrigel growth factor reduced (8.7 *µ*g/cm2, Corning, 356230) coated plates at 37 °C with controlled 5% CO2 and passaged every 3–4 days using 0.7 mM EDTA.

### 4.3 Organoid generation

The STEMdiff Cerebral Organoid Kit (StemCell Technologies 08570) was used to generate optimised Cerebral Organoids (COs). Single cell suspension was obtained through resuspension of stem cells in Accutase (Sigma-Aldrich, A6964). 500 Cells were seeded per well in U-bottom ultralow attachment 96 well plates (Corning, CLS7007) with EB media supplemented with 50 *µ*M ROCK inhibitor Y27632 (Millipore, SCM075). At day 5 EBs were transitioned to neural induction media. At day 7 EBs were transferred to 6 cm dishes and incubated in expansion media with dissolved Matrigel (Corning, 356234) in a dilution of 1 to 50. After another three days of culture, organoids were incubated with Improved Differentiation Media without vitamin A (5% (v/v) DMEM F12 (Thermo Fisher Scientific, 11330032), 50% (v/v) Neurobasal (Invitrogen, 21103049), 1:200 (v/v) N2 supplement (Thermo Fisher Scientific 17502048), 1:50 (v/v) B27-A (Thermo Fisher Scientific, 12587010), 1:100 (v/v) GlutaMAX, 1:200 (v/v) MEMNEAA, 2.5 *µ*g/mL Insulin solution (Sigma-Aldrich, I9278), 50 *µ*M *β*-mercaptoethanol (Life Technologies, 31350-010), 1:100 (v/v) Penicillin-Streptomycin). Cerebral organoids were fixed for 1 hour using 4% PFA at RT at day 15, and washed 3 times in PBS for 30 minutes at RT.

### 4.4 Expansion microscopy

After fixation samples were incubated with 0.1% PBS-Triton X-100 for 2 hours and incubated overnight in a solution of 1.4% FA (28906, Fischer Scientific) and 2% Acrylamide (A9099, Sigma) at 37° C. Samples were incubated at 4° C with monomer solution (19% Sodium Acrylate (408220, Sigma), 1% Acrylamide, 0.1% BIS (J66710.30, Thermo) in 1X PBS). Samples were then placed in a humidified chamber, embedded in monomer solution supplemented with TEMED (411019, Sigma) and APS (A3678, Sigma) to a final concentration of 0.5% and transferred to 37° C for 1.5 hours. The organoid containing gels were then incubated with denaturation buffer (200 mM SDS (30018385, Fischer Scientific), 200 mM NaCl (3624-01, J. T. Baker), 50 mM Tris (30018880,Fischer Scientific), pH 9) for 15 minutes at RT followed by 1 hour and 30 minutes at 95° C. Gels were then subject to a first round of expansion by incubating three times in ddH2O for 30 minutes. The gel was then shrunk by washing 3 times with PBS for 15 minutes. Gels were incubated with primary antibodies diluted in PBS-BSA 2% for 48 hours. Afterwards, they were subject to 3 washes in PBS-Tween 0.1% (P7949, Sigma) and incubated with secondary antibodies and hoescht33342 in PBS-BSA 2% for 48 hours. Once more gels were washed in PBS-Tween 0.1% and subject to a final round of expansion by incubating three times in ddH2O for 30 minutes.

### 4.5 Sample labelling

Primary antibodies used were mouse anti-*α*-tubulin (T9026, Sigma, 1:250) and rabbit anti-NUP96/98 (12329-1-AP, Proteintech, 1:300) and anti-Sox2 (AB5603, Millipore, 1:200). Secondary antibodies were Alexa Fluor 488 donkey anti-mouse IgG (A21202, Thermo Fisher Scientific, 1:250) and Alexa Fluor 568 donkey anti-rabbit IgG (A10042, Thermo Fisher Scientific, 1:250). Nuclear counterstaining was performed using Hoechst 33342 (H3570, Thermo Fisher Scientific, 1:500). Atto 488 NHS ester (41698, Sigma) was used for pan-staining at a concentration of 100 *µ*m in sodium bicarbonate buffer pH 8.3.

### 4.6 Imaging

All imaging was performed using an inverted Nikon CSU-W1, spinning disk. The water dipping objective of choice was the CFI Apo NIR 60X W/1 NA, which has a working distance of 2.8 mm. The Attofluor Cell Chamber (A7816, ThermoFischer Scientific) was fitted with FEP film (RS Components Ltd, 5363996), cut to 1-inch diameter circles to fit using a circular punch (Ezydka Circle Hole Punch, Amazon UK, ASIN: B0D1V931N3). A 0.5-inch punch from the same set was used to cut the FEP film that rests on the objective lens.

## Acknowledgment

The authors gratefully acknowledge the Light Microscopy Facility of the MRC LMB for access to the Yokogawa W1 spinning disk microscope used in this work.

## Data availability

Data underlying the results presented in this article are available from the authors on request.

## Supplementary Information

**Supplementary Figure 1:**
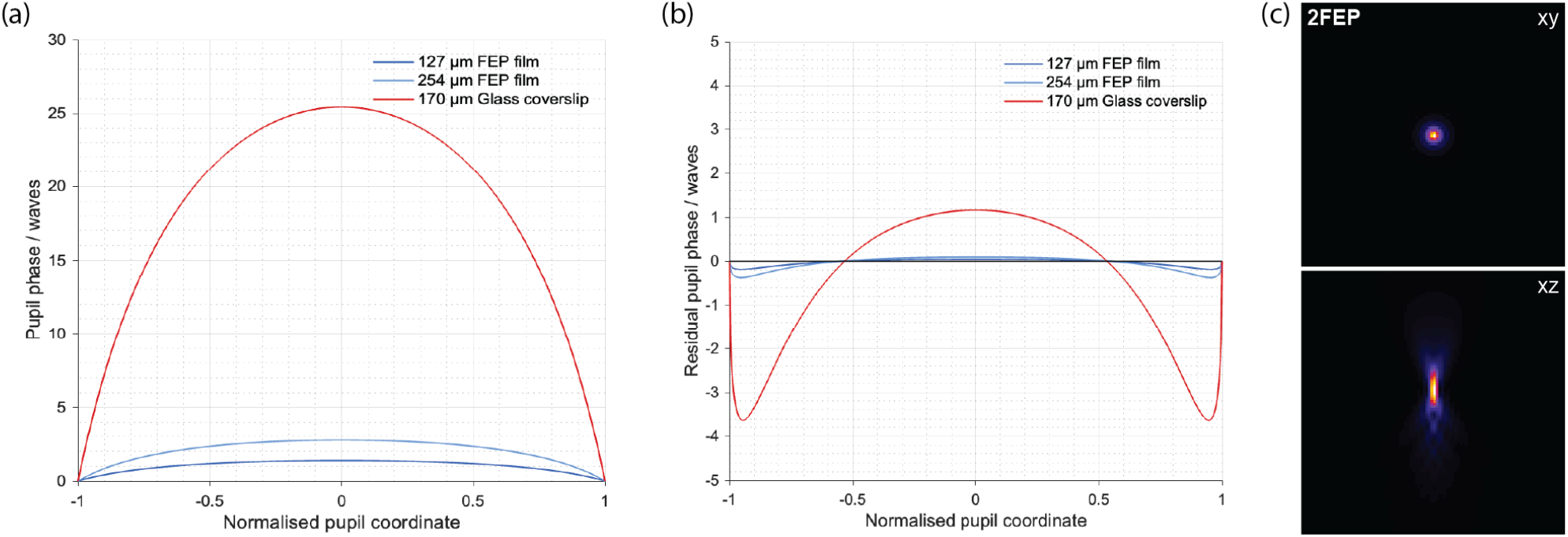
As in Figure 1, but including a third condition with two stacked 127 *µ*m FEP films. (a) Simulated phase aberrations remain dominated by full-defocus. (b) Residual aberrations after full-defocus correction show that both single and double FEP films are effectively corrected, whereas glass retains substantial residual error. (c) Simulated 3D PSF confirms that the use of two FEP films still yields near-ideal PSFs.

**Supplementary Figure 2:**
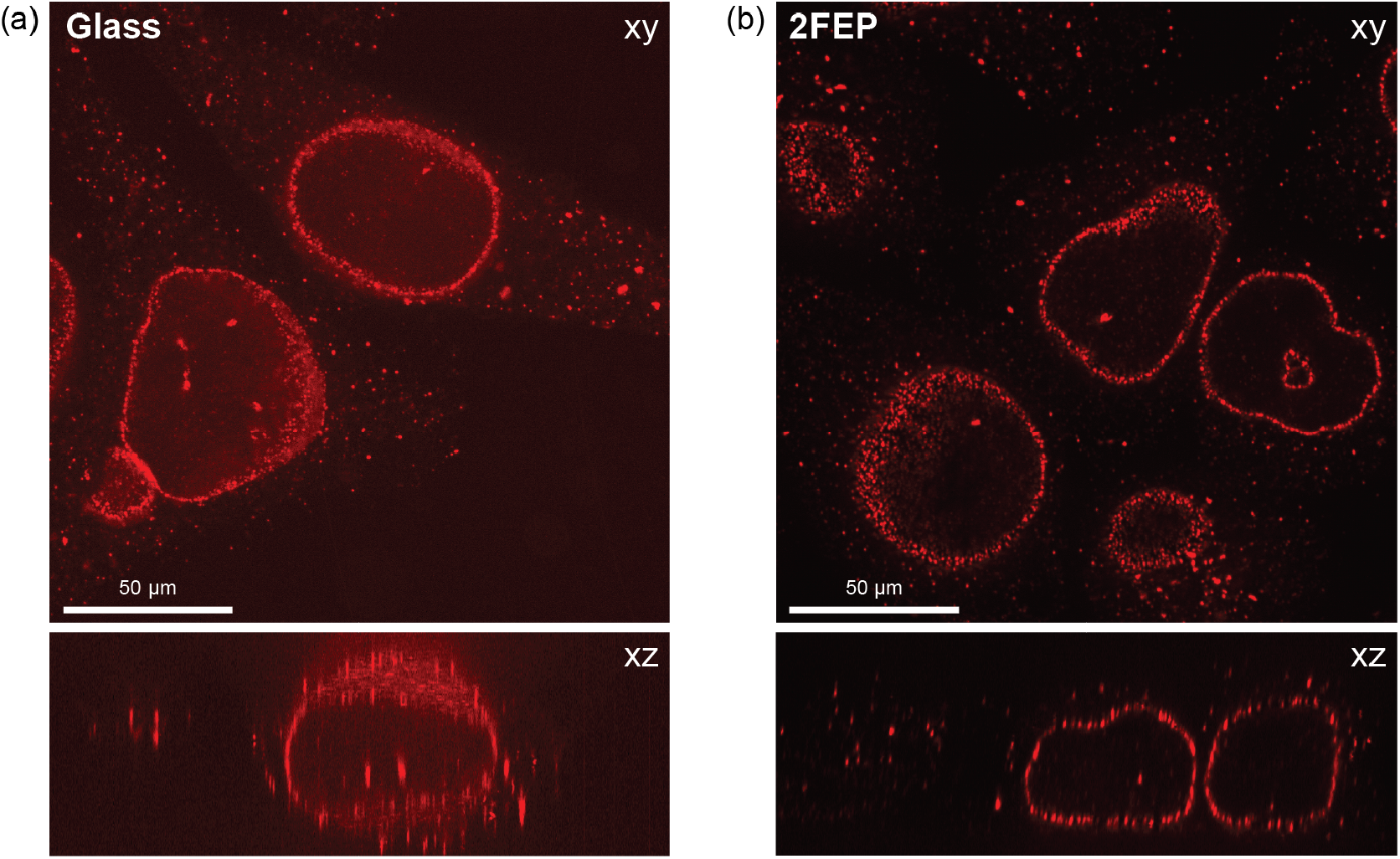
Comparison of imaging 4×-expanded HeLa cells labeled for the nuclear pore complex components NUP96 and NUP98 using (a) a glass coverslip and (b) the proposed 2FEP strategy. Imaging with glass results in strong aberrations and distorted structures, whereas the 2FEP configuration preserves well-resolved nuclear pore organisation.

**Supplementary Figure 3:**
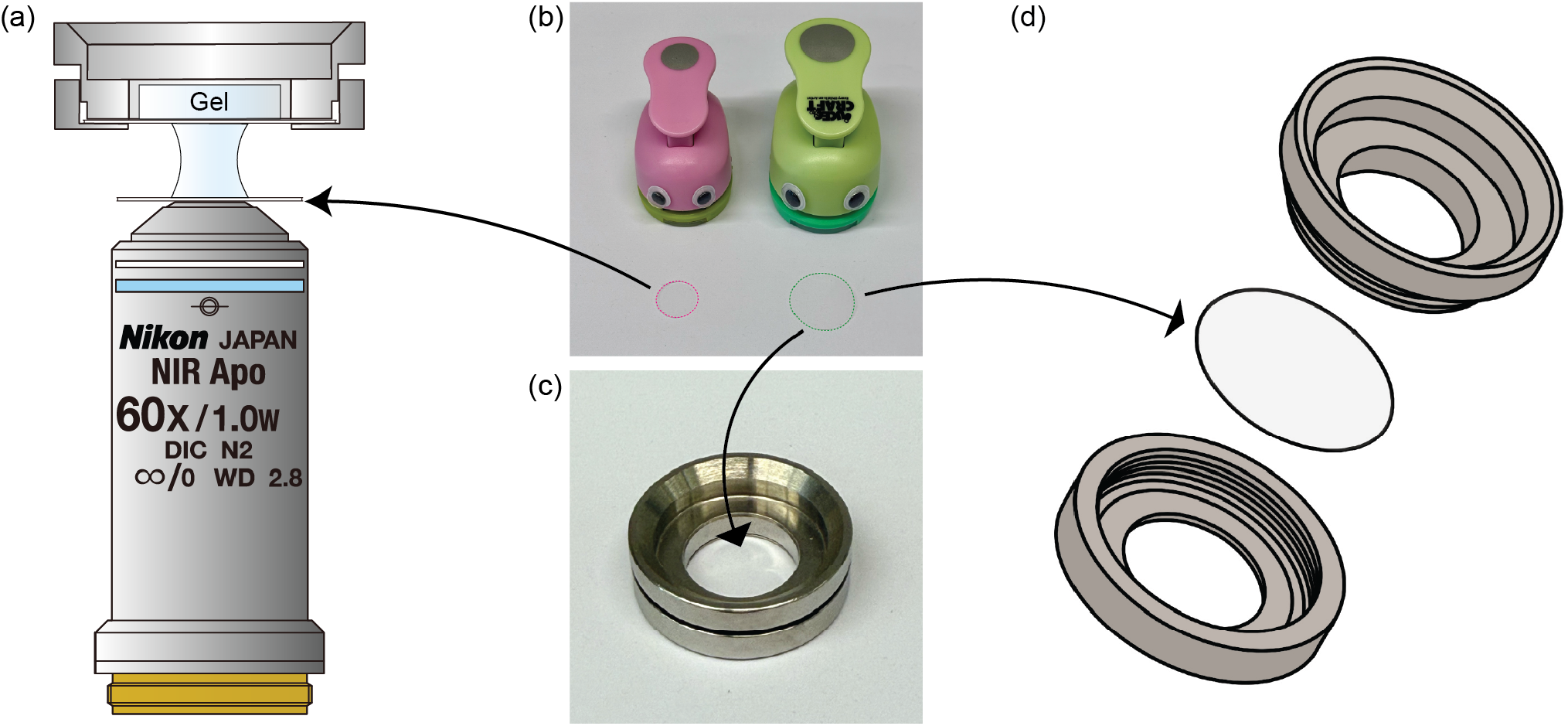
(a) Schematic of imaging framework. (b) FEP punches: 1-inch (green) and 0.5-inch (pink). (c) Attofluor chamber. (d) Exploded view of dish with integrated FEP coverslip.

